# Transient N-6-methyladensosine Transcriptome sequencing reveals a regulatory role of m6A in splicing efficiency

**DOI:** 10.1101/242966

**Authors:** Annita Louloupi, Evgenia Ntini, Thomas Conrad, Ulf Andersson Ørom

## Abstract

Splicing efficiency varies among transcripts, and tight control of splicing kinetics is crucial for coordinated gene expression. N-6-methyladenosine (m6A) is the most abundant RNA modification and is involved in regulation of RNA biogenesis and function. The impact of m6A on the regulation of RNA splicing kinetics has not been investigated. Here, we provide the first time-resolved high-resolution assessment of m6A on nascent RNA transcripts and unveil its importance for the control of RNA splicing kinetics. We identify that early co-transcriptional m6A deposition near splice junctions promotes fast splicing, while m6A modification of introns is associated with long, slowly processed introns and alternative splicing events. In conclusion, by directly comparing the processing dynamics of individual transcripts in the methylated versus unmethylated state on a transcriptome-wide scale we show that early m6A deposition marks transcripts for a fast-track processing.

## Introduction

Splicing of RNA often occurs simultaneously with transcription(Bentley, 2014). Regulation of splicing is essential for functional gene expression and efficient cellular responses to environmental changes. The RNA-binding proteins and *cis*-regulatory elements involved in splicing regulation have been extensively studied and characterized (Wahl and Luhrmann, 2015a, b, c). The RNA nucleotide code is supplemented by more than a hundred chemical modifications, greatly extending the functionality and information content of RNA (Dominissini et al., 2012; Harcourt et al., 2017; Meyer et al., 2012). N-6-methyladenosine (m6A) is the most abundant internal modification in mRNA and non-coding RNA (Dominissini et al., 2012; Meyer et al., 2012; Schwartz et al., 2014). Mapping the occurrence of m6A on steady-state mRNA was accomplished with the development of methyl-RNA immunoprecipitation and sequencing (MeRIP-Seq). This technique uses fragmented RNA as input for m6A immunoprecipitation with an m6A-specific antibody, enabling the identification of ~200 nt long m6A peaks (Dominissini et al., 2012; Meyer et al., 2012). It is has been shown that m6A deposition on steady-state mRNA occurs mostly within the 3’ untranslated region (3’ UTR), around stop codons and within long internal exons (Dominissini et al., 2012; Meyer et al., 2012; Schwartz et al., 2014). Recently, it was reported that m6A methylation is deposited soon after RNA synthesis, with the vast majority of m6A addition observed mostly in exonic sequences when studying chromatin-associated RNA obtained by cellular fractionation (Ke et al., 2017).

m6A is deposited by a protein complex comprised by the methyltransferase-like 3 and 14 (MΕΤΤL3, MΕΤΤL14), Wilms’ tumor 1-associating protein (WTAP), and the Virilizer homolog (KIAA1429) (Liu et al., 2014; Ping et al., 2014; Schwartz et al., 2014). MΕΤΤL3 is the catalytically active subunit, while MΕΤΤL14 plays a structural role critical for substrate recognition, enhancing MΕΤΤL3 activity (Wang et al., 2016a; Wang et al., 2016b). Early studies have demonstrated that adenosine methylation frequently occurs within a subset of RRA*CH consensus sites (R, purine; A*, methylatable A; H, non-guanine base) (Narayan and Rottman, 1988). Fat mass and obesity-associated (FTO) and AlkB homolog 5 (ALKBH5) proteins have the ability to remove adenosine methylation through a multi-step intermediated process including hydromethyladenosine (hm6A) and N6-formylasenosine (f6A) (Fu et al., 2013; Jia et al., 2011; Zheng et al.). Interestingly, FTO and ALKBH5 do not exhibit strict sequence requirements for substrate specificity, and a consensus motif is not crucial for selectivity (Zou et al., 2016).

The function of m6A has been studied in numerous mRNA processes including splicing, RNA degradation and translation (Bartosovic et al., 2017; Ke et al., 2017; Meyer et al., 2015; Wang et al., 2014; Xiao et al., 2016). These pathways are mediated in part by members of the YTH-domain protein family called m6A readers which recognize and bind specifically on m6A sites (Xiao et al., 2016; Xu et al., 2014). Regarding mRNA splicing, YTHDC1 promotes exon inclusion of targeted mRNAs by facilitating binding of SRSF3 and blocking SRSF10 binding to RNA (Xiao et al., 2016). Furthermore, the presence of m6A can affect the RNA structure and increase the accessibility of adjacent RNA sequences for the heterogeneous nuclear ribonucleoproteins HNRNPG and HNRNPC, with an effect on splicing (Liu et al., 2015; Liu et al., 2017). These *trans*-acting RNA-binding proteins together with small nuclear RNAs (snRNAs) recognize and bind pre-mRNA at splice-junctions. Together with *cis* regulatory Exonic Splicing Enhancers (ESEs) these elements orchestrate splicing of the nascent RNA in a coordinated manner (Keene, 2007; Wahl and Luhrmann, 2015b). Although many findings indicate a role of m6A in splicing, a direct link between RNA splicing dynamics and m6A deposition has not been shown. Here, by developing and applying TNT-seq (Transient N-6-methyladensosine Transcriptome sequencing) and qTNTchase-seq (quantitative TNT pulse-chase sequencing) we show that m6A modifications deposited early/co-transcriptionally near splice junctions (5’ and 3’ SJ) positively affect RNA splicing kinetics. Furthermore, we identify intronic m6A sites that are connected with slow processing kinetics and alternative splicing events. Our results strongly support a functional role for m6A in the regulation of splicing, moving yet another step closer to unravelling the multiple facets of epitranscriptomic gene regulation.

## Results

### TNT-seq reveals m6A deposition on newly transcribed RNA

We established TNT-seq, a technique to identify m6A on nascent RNA, enabling us to study the deposition of m6A on short-lived RNA processing intermediates. In brief, we applied MeRIP-Seq on metabolically labeled transcripts that are produced within a 15 minutes window of active transcription (Figure 1A) (Dominissini et al., 2012; Meyer et al., 2012). We chose to use bromouridine (BrU) for labeling of nascent RNA, since it can be incubated in cell media for hours without any toxic effect, ideally suited for *in vivo* studies (Paulsen et al., 2013). Following a 15 minutes BrU-pulse, cells were collected; isolated RNA was heat-fragmented to ~100 nt length and labeled RNA was purified with a BrU-specific antibody (Figure 1A). BrU-labeled RNA was subsequently eluted via BrU competition, to reduce background from contaminating unlabeled RNA, and the eluate was then subjected to imuunoprecipitation with an m6A-specific antibody to enrich for methylated RNA fragments. The BrU-labeled nascent RNA (BrU-RNA Input) and the m6A enriched RNA fragments (BrU-m6A-RNA IP eluate) were then subjected to deep sequencing to identify positions of m6A on nascent RNA (Figure 1A). We detect localized enrichment of m6A deposition at start and stop codons as well as at 5’ and 3’ SJs as a reproducible profile from independent replicates (Supplementary Figure 1A), suggesting a robust experimental pipeline (genome-wide m6A signal correlation = 0.58, see Methods under ‘RNA sequencing and data analysis’). m6A peaks were identified using a published pipeline (Ke et al., 2015; Ke et al., 2017) (See Methods under ‘m6A peak calling’). We find that the majority of early m6A peaks (57 %) reside within intronic sequences, 22 % in coding sequences (CDS), 5 % in 5’ UTRs and 9% in 3’ UTRs (Figure 1B). We obtain a similar m6A peak distribution by calling m6A peaks using MACS2 as in (Dominissini et al., 2013) (not shown). To compare m6A peak distribution in newly transcribed RNA with steady-state mRNA we reanalyzed MeRIP-Seq data from (Schwartz et al., 2014) and called m6A peaks using the same pipeline. The majority of steady-state mRNA m6A peaks reside in the CDS (52 %), 3’ UTR (28 %) and 5’ UTR (12 %), while only a minor fraction (4 %) is intronic (Figure 1C). To validate the authenticity and context of early m6A sites we assessed the presence of the DRACH m6A consensus motif by performing a *de novo* motif search with HOMER (Heinz et al., 2010) in the regions +/-150 nt around the peak summit of best scoring peaks (score > 20, n= 5651) or in randomly generated 300 nt genomic intervals (See Methods under ‘*De novo* Motif Search’). This analysis identified the presence of a DGACH motif with a positional enrichment around the peak summits, in particular for exonic peaks (Supplementary Figure 1B). In addition, by *de novo* motif search we identify three additional motifs, sharing an SAG core, with a strong positional enrichment around the peak summit, especially for intronic peaks (Supplementary Figure 1B). Regarding positional enrichment, almost half (45 %) of CDS-associated nascent m6A peaks reside within a 100 nt exonic window from the 5’ SJ (i.e. within 100 nt upstream of the 5’ SJ), and about one fifth (~18 %) are within 100 nt downstream of the 3’SJ (Figure 1B). In data from steady-state mRNA only 17 and 11 per cent of the CDS peaks are within 100 nt upstream of the 5’ SJ and within 100 nt downstream of the 3’ SJ respectively (Figure 1C), suggesting a transient functional role of early m6A deposition.

**Figure 1:**
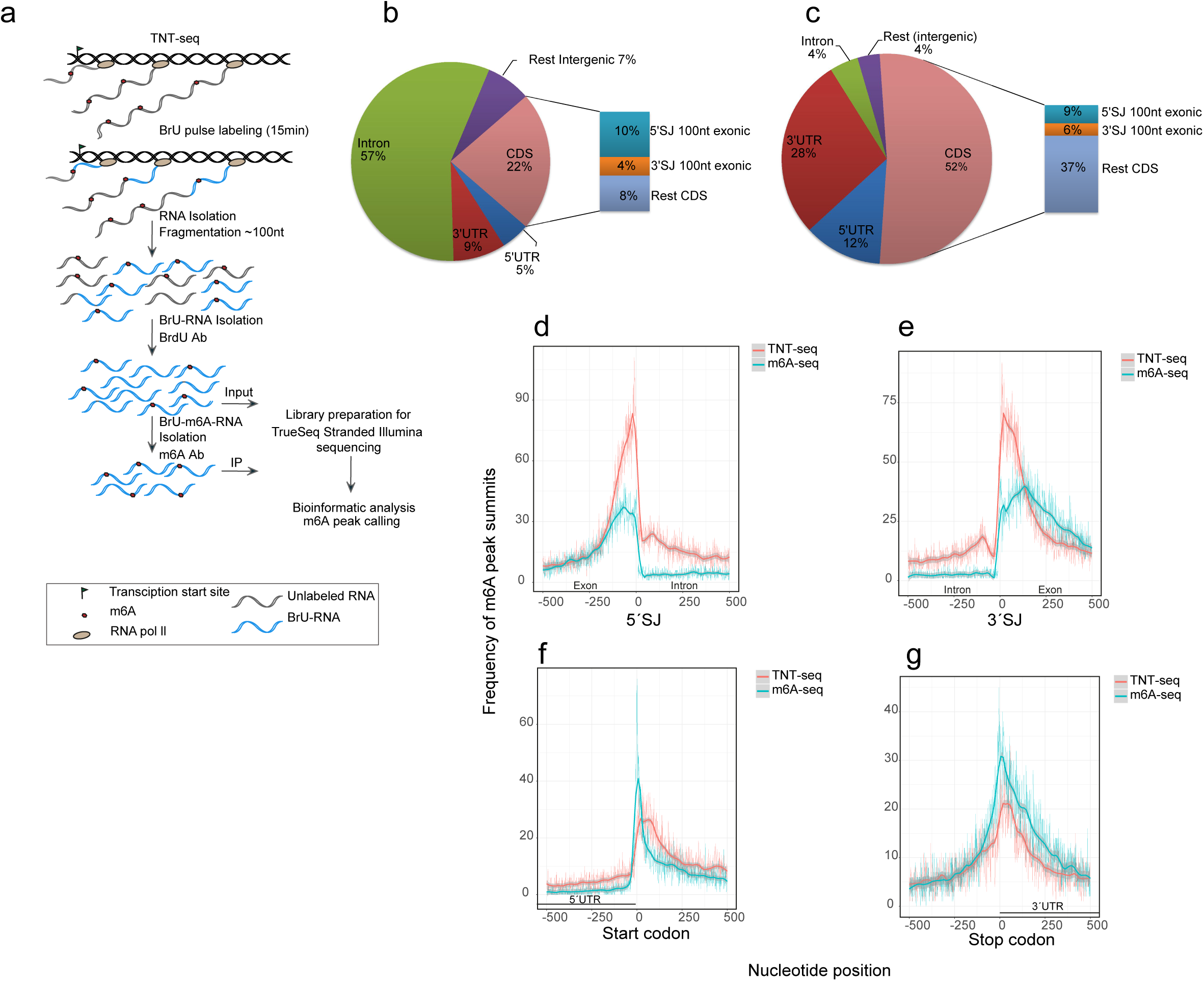
TNT-seq reveals m6A deposition on newly transcribed RNA. **(a)** Schematic representation of the TNT-seq protocol. **(b-c)** m6A peak distribution in **(b)** newly transcribed RNA and **(c)** mRNA(Schwartz et al., 2014). **(d-g)** Distribution (frequency) of the distances of m6A peak summits to the closest given anchor point **(d)** 5’SJ, **(e)** 3’SJ, **(f)** Start codon and **(g)** Stop codon for newly transcribed RNA (TNT-Seq) and mRNA (m6A-Seq(Schwartz et al., 2014)). Distance of a peak summit to the closest given anchor point was calculated with bedtools closest.

We then analyzed the positional distribution of m6A peak summits around 5’ SJs, 3’ SJs, and start- and stop-codon anchor points for both newly transcribed and steady-state mRNA (Figure 1D-G). Early m6A deposition predominates at and in close proximity to splice junctions (Figure 1D-E), whereas the picture is inversed around start- and stop-codons, with a relatively greater number of peaks in steady-state mRNA (Figure 1F-G). This finding led us to examine whether early m6A deposition in close proximity to SJs has an impact on splicing of RNA.

### m6A signatures separate distinct intron classes

To determine the splicing kinetics of newly transcribed RNA, we used BrU-pulse-chase sequencing as described (Louloupi et al., 2017; Paulsen et al., 2013). In brief, cells were labeled with a 15 minutes BrU pulse and chased for 0, 15, 30 and 60 minutes followed by RNA purification. We calculate the splicing index value θ (Mukherjee et al., 2017) (Supplementary Figure 2A) to determine splicing efficiency across all time points for introns that have at least 5 reads coverage on both 5’ and 3’ SJ for all included RNA sequencing libraries (four time points of BrU-pulse-chase-seq and the three Input samples for TNT-seq), yielding 13,532 introns with an extracted θ value ranging from 0 (unspliced) to 1 (fully spliced) (Methods). As expected, the cumulative distribution of the splicing index at 0 min, representing nascent RNA, precedes that of the steady-state chromatin-associated RNA (Conrad et al., 2014) indicating that pre-mRNA is efficiently captured by the assay (Figure 2A). Using k-means clustering with k = 3 we called three clusters of distinct splicing efficiency dynamics (SED) representing 4,882 fast, 5,702 medium, and 2,948 slowly processed introns (Figure 2B and Supplementary Figure 2B-D) (for the definition of SED see Methods under ‘Splicing kinetics and predictive models’). Snapshots from the UCSC genome browser for three representative cases are depicted in Supplementary Figure 2E. To examine how early m6A deposition varies with different processing efficiencies, we plotted the average m6A signal per nucleotide position around 5’ and 3’ SJ (Figure 2C-D) and within length-binned introns for the three groups (Supplementary Figure 3A-C). Strikingly, we find that fast processed introns show greater m6A deposition at SJs with an overall positive relationship between m6A deposited at 5’ and 3’ SJ exonic boundaries and processing efficiency (Figure 2C-D and Supplementary Figure 3A, 3C). By plotting the average frequency of m6A peak summits per nucleotide position (instead of the average m6A signal) for the three subgroups, we reach the same conclusion (Supplementary Figure 3D, 3F). In contrast, slowly processed introns are associated with increased m6A deposition within the intron (Supplementary Figure 3B, 3E).

**Figure 2:**
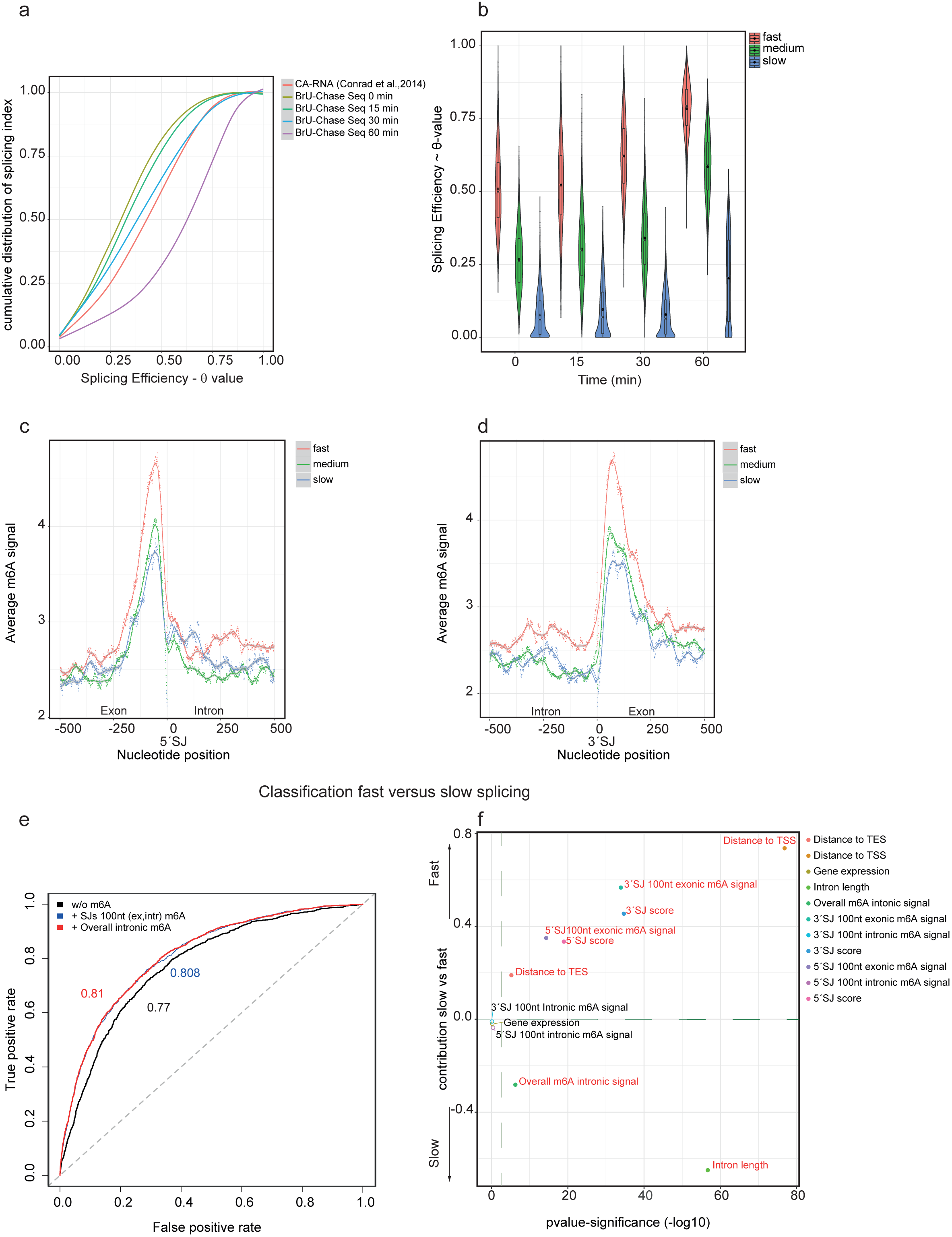
m6A deposition at nascent RNA predicts splicing efficiency dynamics. **(a)** Cumulative distribution of the splicing efficiency index from chromatin-associated RNA seq(Conrad et al., 2014), BrU Chase Seq 0 min, BrU Chase Seq 15min, BrU Chase Seq 30min and BrU Chase Seq 60 min **(b)** Violin plot representing the density of the distribution of the splicing efficiency index (θ value) with embedded box and whisker plots for introns grouped on the basis of differential splicing kinetics (see also Supplementary Figure 2A-D). **(c-d)** Average m6A signal per nucleotide position in the window +/- 500 nt around (**c**) 5’SJ and 3’SJ (**d**) for 13,532 introns. **(e)** Average receiver operating characteristics (ROC) curve for discrimination of fast versus slow introns including all characteristics and excluding m6A. The respective Area Under the Curve (AUC number) is indicated. **(f)** Plot depicting contribution of each feature to the model fit of fast versus slow processing calculated as the coefficients from the binary logistic regression with the associated estimated significance (-log10 p-value). The features with p-value <0.001 are red-colored.

### m6A deposition at nascent RNA predicts splicing efficiency dynamics

To further investigate the impact of m6A deposition on nascent RNA in shaping the splicing efficiency dynamics we used several features in a logistic regression model fit to predict fast versus slowly processed introns (Methods) (Figure 2E-F). We find that inclusion of the m6A at SJs as an additional parameter improves the predictive power of the model (Figure 2E) with the m6A contribution in predicting fast processing being comparable to other previously shown features (Mukherjee et al., 2017), such as the 5’ and 3’ SJ sequence scores and distance to TSS/TES (Figure 2F). In contrast, the overall intronic internal m6A signal and intron length are significantly associated with slow processing (Figure 2F). To complement this analysis we further employed linear regression to predict splicing efficiency dynamics (SED) as a continuous value (Supplementary Figure 4). Again here, introducing the m6A at SJs improves the correlation between predicted and measured SED (Supplementary Figure 4A-C) further confirming the impact of early m6A deposition on the efficiency of RNA processing.

### Internal intronic m6A deposition associates with alternative splicing

Slow pre-mRNA processing has been shown to favor the occurrence of alternative splicing, *i*.*e*. exon-skipping (Mukherjee et al., 2017). We assessed alternative versus constitutive splicing by extracting the intron-centric ψ value as in (Mukherjee et al., 2017) (Supplementary Figure 2A and Methods). Our data further support that alternative splicing events are significantly enriched in slowly processed introns (odds ratio 3.84, Fisher’s exact test p-value < 2.2e-16) (Figure 3A). We next asked whether intronic m6A deposition could affect alternative splicing. We find that intronic m6A peaks associate with upstream or downstream exon-skipping about two times more often than expected by random chance (odds ratio 1.7, Fisher’s exact test p-value < 2.2e-16), indicating that internal intronic m6A deposition is significantly enriched in alternative splicing events. In concurrence, we find that the average m6A signal is greater along alternative versus constitutively spliced introns (Figure 3C). As expected, the average m6A signal is greater at constitutive versus alternatively spliced SJ exonic boundaries (Figure 3B, 3D). In the prediction of alternative versus constitutive splicing, the overall intronic m6A, along with the physical characteristic of intron length, are significant contributors in determining alternative splicing (Figure 3E). In contrast, m6A at SJ exonic boundaries and strong splice site consensus sequences (SJ score) ensure constitutive splicing (Figure 3E). As in predicting fast versus slow processing (Figure 2E), inclusion of m6A also improves the predictive power of the model fit of constitutive versus alternative splicing (Figure 3F), once more highlighting the impact of m6A deposition on nascent RNA in shaping splicing efficiency.

**Figure 3:**
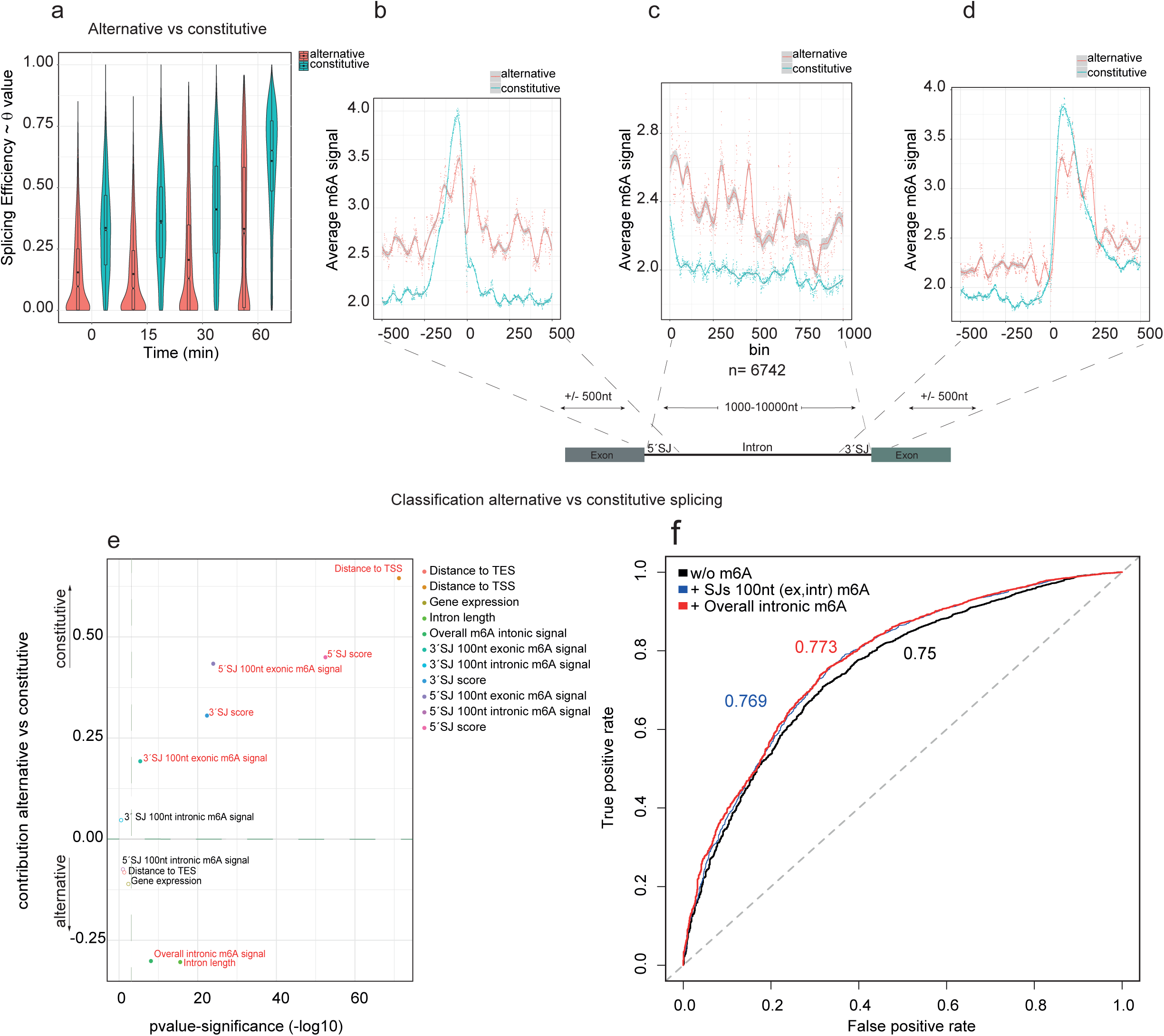
Internal intronic m6A deposition associates with alternative splicing. **(a)** Violin plots representing the density of the distribution (with embedded box-and-whiskers plots) of the splicing efficiency index (θ value) for introns classified as either constitutive or alternative (i.e. showing adjacent exon skipping) spliced, extracted from all pulse-chase time points. **(b-d)** Average m6A signal per nucleotide position in a +/-500 nt window around **(b)** the 5’ SJ and **(d)** 3’ SJ, and per bin **(c)** of 6,742 introns with 1,000–10,000 nt length. The average m6A signal is extracted separately for the two subgroups, constitutive and alternative. The lines represent loess curve fitting (local polynomial regression) with the 95% confidence interval grey shaded. **(e)** Plot depicting the contribution of each feature to alternative versus constitutive splicing, calculated as the coefficients of the binary logistic regression fit, with associated estimated significance (-log10 p-value). Features with p<0.001 are red colored. **(f)** Average receiver-operating characteristic curve (ROC) for the logistic regression prediction of the alternative versus constitutive splicing using all features, with and without m6A data. The respective Area Under the Curve (AUC number) is indicated.

### qTNTchase-seq identifies m6A-marked fast-track RNAs

After identifying intron classes with distinct m6A signatures and processing kinetics, we wanted to assess the direct impact of m6A modifications at the individual transcript level. To clearly separate directly m6A-mediated from sequence specific effects on RNA processing, we developed and applied qTNTchase-seq (quantitative TNT pulse-chase sequencing). Here, labeled RNA was isolated at 0 and 30 min chase after the BrU-pulse and immunoprecipitated with an m6A-specific antibody without prior fragmentation to maintain transcript level information (Figure 4A). Both supernatant (m6A negative transcripts) and eluate (m6A positive transcripts) were sequenced for each time-point to obtain quantitative information. We performed two biological replicates of qTNTchase-seq and calculated the m6A levels per transcript according to (Molinie et al., 2016) (Methods). On a transcriptome-wide scale we observe a strong concordance of m6A levels between the two biological replicates, irrespective if only the top 25% expressed transcripts or all transcripts with non-zero coverage are included in the analysis (for 0 minutes Pearson *r* = 0.89 p value < 2.2e-16 and for 30 min Pearson *r* = 0.91 p value < 2.2e-16) (Supplementary Figure 5A-B). When comparing m6A levels between 0 min and 30 min chase we do not observe any significant differences indicating that overall m6A modification levels on transcripts remain the same for at least ~45 minutes after transcription (Supplementary 5C). We then analyzed splicing efficiency on the transcript level by extracting the transcript splicing index (Methods under ‘Transcript m6A level and splicing index’) and compared this for methylated versus non-methylated transcripts at 0 min and 30 min separately. Within the pulse, corresponding to a 15 minute window of transcription, methylated transcripts show significantly higher splicing efficiency than non-methylated transcripts (Figure 4B), further supporting the role of the early m6A deposition in enhancing processing efficiency. In addition, by measuring the splicing efficiency dynamics (SED) at the transcript level from 0 to 30 minutes chase, we find that methylated transcripts show on average significantly greater processing than unmethylated transcripts (two tailed paired *t-test* p-value < 2.2e-16) (Figure 4C). Importantly, processing appears significantly enhanced for the same individual transcripts in the methylated compared to the unmethylated state; ~76% of the transcripts show gain of SED in the methylated versus unmethylated state revealing a direct and sequence independent impact of m6A on processing kinetics (Figure 4D). We further examined locally the splicing efficiency for the dataset of the 13,532 filtered introns. We find, that ~14% have significantly higher splicing efficiency in the m6A positive than in the m6A negative transcripts and show a 1.26 fold enrichment over random chance to possess an m6A peak in the 5’ SJ 250 nt exonic boundary (odds ratio 1.265, Fisher’s exact test p-value 0.0006745). In addition, individual intron loci show on average significantly higher SED in methylated versus non-methylated transcripts (two tailed paired *t-test* p-value < 2.2e-16) (Supplementary figure 5D).

**Figure 4:**
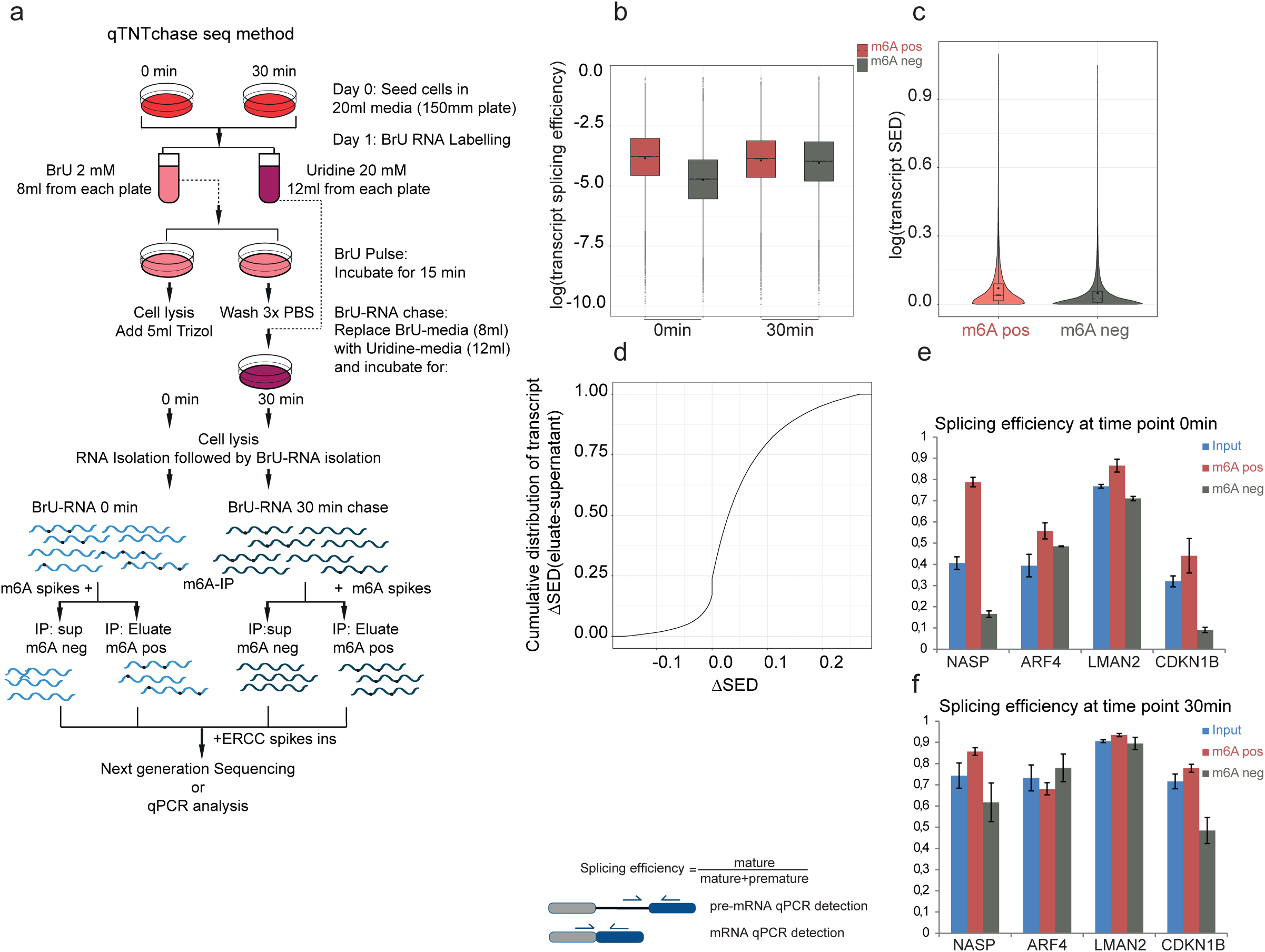
qTNTchase-seq identifies m6A-marked fast-track RNA. **(a)** Schematic description of the qTNTchase-seq method. **(b)** Box plot representing the overall splicing efficiency of methylated (m6A positive) versus non methylated (m6A negative) transcripts at time points 0 min and 30 min. **(c)** Violin plots showing distribution of the transcript SED in m6A positive and m6A negative fractions. (**d**) Cumulative distribution of transcript SED differences between methylated and unmethylated state (∆SED = SED m6A-positive – SED m6A-negative) **(e, f)** qPCR analysis to measure the local intronic splicing efficiency of methylated versus non-methylated transcripts for **(e)** 0 minutes and **(f)** 30 minutes.

We next analyzed the splicing kinetics of four candidate splice junctions that possess at least one m6A peak (+/-250nt) by qPCR on qTNTchase-seq RNA. We determined splicing efficiency as the ratio of the spliced signal over total (spliced + unspliced) signal. Strikingly, at time point 0 min, methylated transcripts show higher splicing efficiency compared to their unmethylated counterparts that share the same nucleotide sequence (Figure 4E-F). This result was also confirmed with RT-PCR analysis where the fragments corresponding to spliced and unspliced transcripts were analyzed on an agarose gel, confirming the positive effect of m6A on RNA splicing (Supplementary Figure 6A-D).

### Splicing factors coincide with m6A deposition

Due to the prominent presence of the early m6A mark at the SJ exonic boundaries and since both the transcriptome-wide and locus-specific experimental data support a positive role of early m6A deposition in regulating processing efficiency, we sought to investigate how this functionality is mediated. For this we analyzed available CLIP-data for SRSF factors with an established role in splicing (Xiao et al., 2016). We find that both SRSF3 and SRSF10 in particular show a high probability to have an m6A peak summit in close proximity (< 250 nt) (Figure 5A-B), with SRSF10 showing relatively greater affinity (Figure 5C). In addition, the SAG core that we identify by *de novo* motif search in early m6A peaks (Supplementary Figure 1B) is reminiscent of the SRSF binding site motifs (Ajiro et al., 2016b; Xiao et al., 2016). Further, both SRSF3 and SFRF10 have been shown to bind near m6A (Xiao et al., 2016). In particular, while SRSF3 binding can be synergistically augmented through interaction with YTHDC1, SRSF10 can independently bind to m6A modified regions (Xiao et al., 2016). In agreement, we find that the ratio of SRSF10/SRSF3 binding is greater at the SJ exonic boundaries for fast processed introns, and internally along within slowly processed introns (Figure 5D-F), in concordance with the respective relative enrichment of early m6A deposition (Figure 2C-D and Supplementary Figure 3). At the same time, the average ratio of SRSF10/SRSF3 binding clearly separates alternative and constitutive spliced introns (Figure 5 G-I), most prominently along length-binned introns (Figure 5H). This result is in agreement with a previous notion suggesting that alternative splicing activity can be antagonistically regulated by SRSF10 versus SRSF3 binding (Xiao et al., 2016). The above data together support that early (co-transcriptional) m6A deposition along intronic sequences could play an early role in shaping the final outcome of alternative splicing activity via resolving the relative recruitment of various splicing-involved regulatory factors with varying m6A affinities.

**Figure 5:**
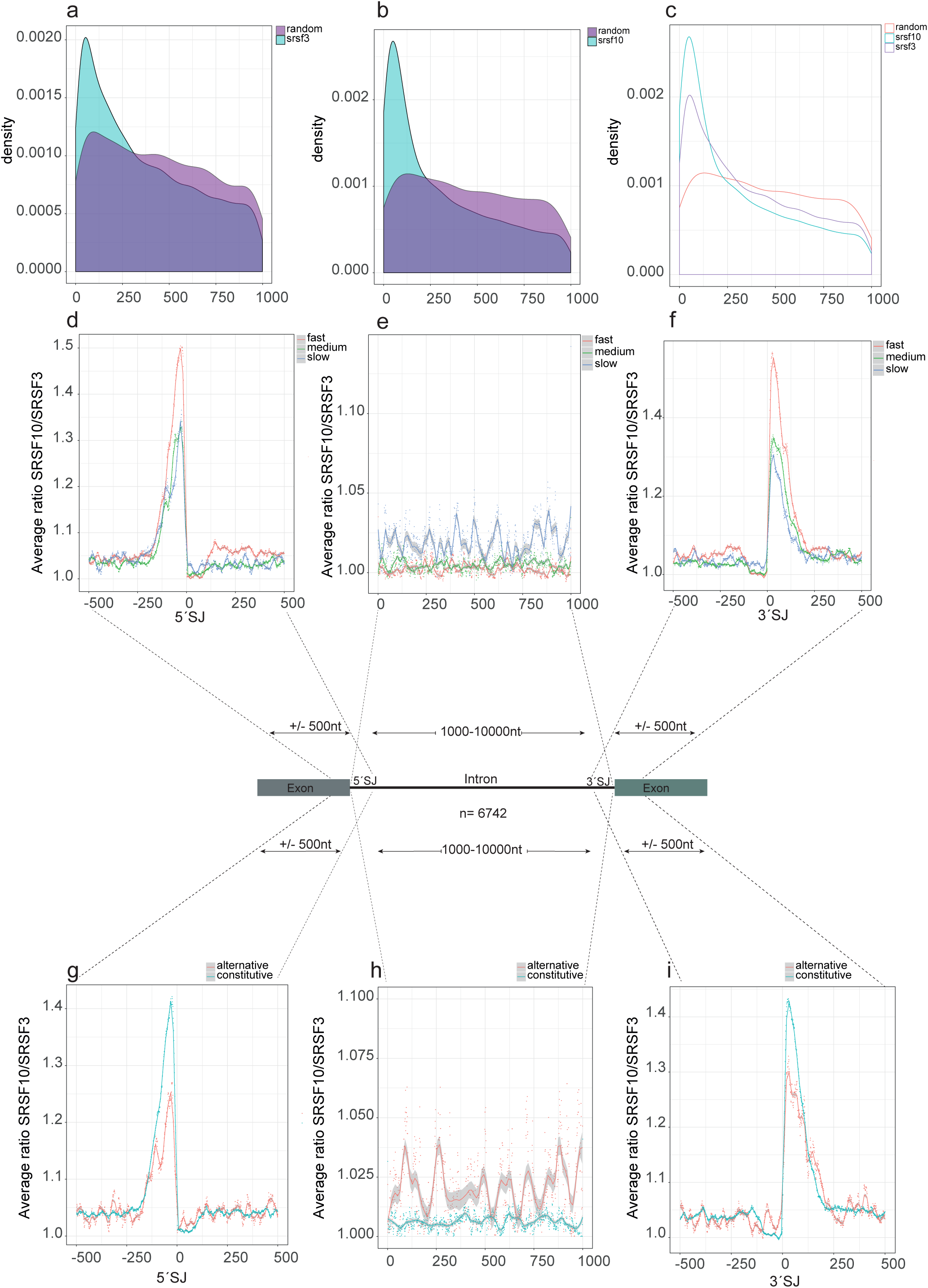
Splicing factors coincide with m6A deposition. **(a-c)** Distribution of the interdistances of factor binding sites to closest m6A peak summit for **(a)** SRSF3 **(b)** SRSF10 and **(c)** overlap. As a control, distance from the midpoint of the respectively same number of randomly generated genomic intervals is also plotted. **(d-f)** Distribution of the average ratio SRSF10/SRSF3 binding, extracted separately for the three subgroups fast/medium/slow per nucleotide position in the window +/-500 nt around the 5’SJ **(d)** and 3’SJ **(f),** or perbin (**e**) for 6,742 length-binned introns (with a length 1,000–10,000 nt). **(g-i)** Same analysis as in (**d-f**) but comparing the average SRSF10/SRSF3 ratio for the two subgroups constitutive versus alternative.

## Discussion

The current study provides the first high-resolution view of the transient, nascent N-6-methyladenosine transcriptome. A recent study from Ke et al.(Ke et al., 2017), reported that m6A is deposited mostly on chromatin-associated pre-mRNAs while observing an enrichment of m6A peaks in exonic regions (Ke et al., 2017). In contrast to this and other studies using steady-state RNA (Dominissini et al., 2012; Meyer et al., 2012; Schwartz et al., 2014), direct assessment of modifications on nascent RNA via TNT-seq reveals that most m6A is initially deposited within intronic sequences, consistent with METTL3-METTL14 PAR-CLIP data showing 29 %-34 % intronic binding sites (Liu et al., 2014).

TNT-seq in conjunction with BrU pulse-chase reveals that the signature of early m6A deposition at splice junctions and within introns is associated with distinct RNA processing kinetics. Most importantly, qTNTchase-seq enabled us to directly compare the processing of individual transcripts in the methylated versus unmethylated state, demonstrating that m6A directly controls splicing kinetics irrespective of the underlying transcript sequence. This finding suggests that m6A serves as a labeling signal that could be recognized by m6A reader proteins to sort methylated transcripts into a fast-track processing dependent on the positional context within the transcript.

Furthermore, our findings reveal that intronic m6A peaks are enriched in introns involved in alternative splicing. Bartosovic et al.(Bartosovic et al., 2017) showed that FTO, an m6A demethylase, binds mostly to introns, mediating m6A removal. FTO knockout causes alternative splicing events with a preference for exon-skipping, suggesting that demethylation of mRNA transcripts promotes exon-inclusion under normal conditions (Bartosovic et al., 2017). Taken together, these findings suggest that intronic m6A marks that are not targeted or not yet removed by FTO mediate exon skipping while introns involved in constitutive splicing show no enrichment in the m6A signal and most probably are targets of FTO (Bartosovic et al., 2017).

In mRNAs, m6A is enriched in the consensus DRACH motif; however not all DRACH motifs are methylated, indicating that the presence of the sequence motif alone is not enough to drive m6A deposition. FTO CLIP data show no significant enrichment of the DRACH motif (Bartosovic et al., 2017) leading us to hypothesize that early m6A intronic deposition is mostly in non-DRACH sequences where FTO can detect and eventually remove the m6A marks. We find a DGACH motif with a positional enrichment around the m6A peak summit, especially for exonic m6A peaks. Using *de novo* motif analysis we identified three additional motifs, sharing an SAG core. These three motifs show strong positional enrichment around the peak summit, especially for intronic peaks, and have higher positional enrichment compared to the DRACH motif. Interestingly, the SAG core is reminiscent of the SRSF binding site consensus.

Recently, Xiao et al. (Xiao et al., 2016) demonstrated that the m6A reader YTHDC1 recruits SRSF3 while competing away SRSF10 and binds to m6A sites promoting exon inclusion. In the absence of YTHDC1 and SRSF3, SRSF10 has the availability to bind to free m6A sites independently, promoting exon skipping. This is also supported by a previous study from Ajiro et al., 2016 showing that SRSF3 knockdown in U2OS cells causes exon skipping events (Ajiro et al., 2016a). The three novel *de novo* found motifs that are enriched in our m6A peaks resemble binding sites of SRSF splicing factors including SRSF3 and SRSF10 (Ajiro et al., 2016b; Xiao et al., 2016). When we calculated the average SRSF10/SRSF3 ratio per nucleotide position for the three subgroups fast, medium, slow and constitutive versus alternatively spliced transcripts we observed a similar distribution and profile to the m6A signal, confirming the competitive binding of SRSF10 versus SRSF3 to m6A regions.

We propose a model (Figure 6), in which the methyltransferase complex co-transcriptionally deposits m6A, probably while being physically bound to RNA PolII (Slobodin et al., 2017). In the case of constitutive splicing, YTHDC1 recruits SRSF3 to SJs that are methylated while FTO binds and demethylates intronic m6A marks. Competition between YTHDC1/SRSF3 and FTO drives the final outcome based on the RNA-binding affinity and protein stoichiometry. Here, features such as 5’ and 3’SJ sequences together with (-100) 5’ SJ and (+100) 3’SJ methylation increase the affinity of splicing factors to promote fast processing kinetics. In the case of alternative splicing, the intronic sequences that are not targeted by FTO are more prone to be bound by SRSF10, thus promoting exon skipping events. Another possibility is that SRSF10, which has high affinity to m6A, binds to methylated introns preventing FTO binding and demethylation of m6A sites. Long introns could potentially have more methylation sites and SRSF10 binding sites, thus are more pronounced to be alternative spliced and slowly processed.

**Figure 6:**
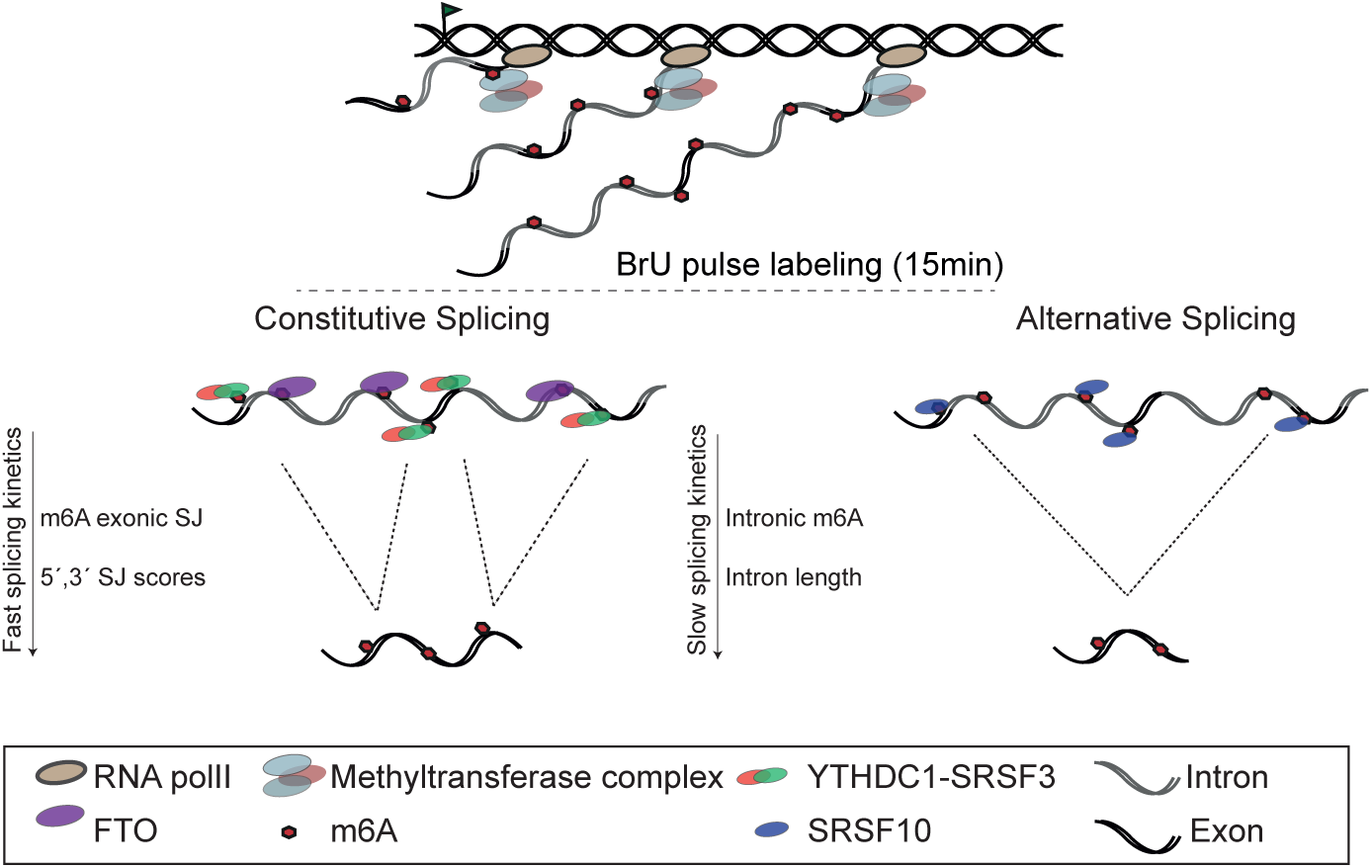
Proposed model scheme. The methyltransferase protein-complex deposits m6A co-transcriptionally while being physically attached to RNA PolII. In the case of constitutive splicing, YTHDC1 recruits SRSF3 to the methylated SJs while FTO is removing the m6A from introns. Here, the m6A presence at SJs together with optimal SJ sequences promotes fast splicing kinetics. In the case of alternative splicing, SRSF10 binds to introns that are not yet or not targeted by FTO promoting exon skipping. Long, methylated introns potentially have more binding sites for SRSF10 thus are more pronounced to be alternative spliced and slowly processed.

The epitranscriptome code is emerging and we are still far from fully understanding it. Many other RNA modifications could contribute to the regulation of RNA processing, thus different combinations of RNA modifications could drive the final outcome. The lack of strong sequence consensus at splice junctions in many introns might be supplemented by the presence of m6A that could eventually attract splicing factors to exert their function. Our study shows that the crucial role of m6A on splicing efficiency dynamics as well as on alternative splicing is positional dependent. m6A deposited in intronic regions sorts transcripts to a slow-track processing pathway and is associated with alternative splicing while, m6A found in splice-junction exonic boundaries, sorts transcripts to a fast-track processing pathway and constitutive splicing.

## METHODS

### Materials and Methods

#### Cell culture and BrU-chase Seq

HEK293 cells were cultured in DMEM growth-medium supplemented with 10% Fetal Bovine Serum (FBS) under normal growth conditions (37°C and 5% CO_2_). The day before bromouridine (BrU) labelling ~2.0 × 10^6 cells were seeded in 100 mm plates with 10ml media, one plate for each time point. Cells were 70–80% confluent before the addition bromouridine (BrU). BrU (-5-Bromouridine cat.no. CAS 957–75–5 Santa Cruz Biotechnology) was added to a final concentration of 2 mM to the media and cells were incubated at normal growth conditions for 15 minutes (pulse). Cells were washed thrice in PBS and either collected directly (0 minutes chase time point) or chased in conditional media supplemented with 20 mM uridine (Sigma cat.no U3750–25G) for 15, 30 and 60 minutes. RNA was purified using TRIzol following manufacturer’s instructions.

In this step we followed the protocol of (Paulsen et al., 2013) with some modifications. 35ul of anti of anti-mouse IgG magnetic Dynabeads (Invitrogen) were transferred to a 1.5ml microfuge Protein Low binding tube and washed 3 times with BrU-IP 1X buffer (0.1% BSA in RNAse free PBS). After the final wash, the beads were resuspended with BrU-IP 1X buffer supplemented with SUPERase• In™ RNase Inhibitor 1:2000 together with BrdU antibody (5μg of antibody per 100 μg RNA). Antibody-beads mixture was incubated for 1hour at room temperature with gentle rotation following 3 washes with 1X BrU-IP. 150 μg RNA was used for each BrU-IP and heated up for 4 minutes at 65°C prior to IP. The same amount of unlabeled total RNA was used as a negative control. 5X BrU-IP (0.5% BSA 5X PBS supplemented with SUPERase• In™ RNase Inhibitor 1:2000) was added to the RNA to have a final concertation of 1X. RNA-antibody-beads mixture was incubated for 90 minutes at room temperature with gentle rotation in a final volume of 800 μl. The beads were washed thrice with 800 μl 1X BrU-IP at room temperature. For all wash steps, with the exception of the elution step, the beads were washed for 5 min rotating then placed on a magnetic rack and the wash buffers were discarded. At the last wash the Protein low binding tubes were replaced with DNA LoBind tubes. For elution 200 μl of Elution buffer (0.1% BSA and 25 mM bromouridine in PBS) were added directly on the beads and the tubes were incubated for 60 minutes with continuous shaking (1100 rpm) at 4 °C. The supernatant (eluate w/o beads) was transferred to a new tube and RNA was precipitated by adding 1/10 volumes of 3M sodium acetate (pH 5.2) and 3–4 volumes of 100% ethanol. RNA was allowed to precipitate at −80 °C overnight. RNA pellet was washed twice with 75% ethanol and resuspended in RNase-free water. RNA quality was analyzed using Agilent 2100 Bioanalyzer with an Agilent RNA 6000 Pico kit according to the manufacturer’s instructions.

#### TNT-seq

For one TNT-seq sample ~ 25 150mm plates were used for BrU labelling. RNA was metabolically labelled with BrU for 15 minutes and RNA was isolated as described above. RNA concentration was adjusted to 2μg/μl with nuclease free water. 18 μl of RNA was added to thin-walled 200µl PCR tube following addition of 2 μl of 10X fragmentation mixture (containing 800 µl of RNase-free water, 100 µl of 1M Tris-HCl pH 7.4 and 100 µl 1M of ZnCl_2_). After vortex and quick spinning the tubes were incubated in 94 °C for 3.5 minutes in a preheated thermal cycler block with the heated lid closed. Tubes were quickly removed from the thermocycler and placed on ice following addition of 2 µl of 0.5 M EDTA. After vortex and quick spin the RNA was collected in a tube to continue with for RNA precipitation using 1/10 volumes of 3 M sodium acetate (pH 5.2), 3–4 volumes of 100% ethanol. RNA was allowed to precipitate at −80 °C overnight. The following day tubes were centrifuged at full speed for 30 minutes at 4 °C. RNA pellet was washed twice with 75% ethanol and resuspended in 400–500 μl of RNase-free water. Validation of post fragmentation size (~100 nt) distribution was analyzed using Agilent 2,100 Bioanalyzer with an Agilent RNA 6,000 Pico kit according to the manufacturer’s instructions. 400 μg-600 μg fragmented BrU labeled total RNA was used for each BrU-IP. BrU-RNA isolation was performed as described above. The BrU-IP recovery was approximately 0.09–0.16% of input. 4.5 μg of BrU fragmented RNA was used as input for the m6A immunoprecipitation. 35 μl of Dynabeads^®^ Protein A (Invitrogen) per sample was transferred to a 1.5 ml microfuge Protein LoBind tube and washed 3 times with 1X m6A-IP (500 mM NaCl, 0.1% NP-40, 10 mM Tris-HCl, pH 7.5). After final wash the beads were resuspend in 800 μl 1X m6A-IP buffer supplemented with SUPERase• In™ RNase Inhibitor 1:1000. 1μg of affinity purified anti-m6A polyclonal antibody (Synaptic Systems) per 2.5 μg BrU-RNA was added to the beads and incubated for 60 minutes at room temperature with gentle rotation. As a negative control, we used Dynabeads^®^ Protein A magnetic beads bound to an irrelevant IgG. Beads were washed 3 times with m6A-IP 1X buffer for 5 min on the rotator. 5 μg of BrU Fragmented RNA was used as input. RNA was heated up for 4 minutes at 65°C. 5X m6A-IP buffer (50 mM Tris-HCl, 750 mM NaCl and 0.5% (vol/vol) Igepal CA-6300 supplemented with SUPERase• In™ RNase Inhibitor) was added to have the RNA in 1X m6A-IP buffer. RNA-antibody-beads mixture was incubated for 2h at 4°C with gentle rotation in a final volume of 0.8ml in Protein low binding tubes. Three washing steps followed using m6A-IP 1X buffer (1^st^ and 2^nd^ wash) and high salt m6A-IP buffer (500 mM NaCl, 0.1% Igepal CA-6,300, 10 mM Tris-HCl, pH 7.5) (3^rd^ wash). For all wash steps, with the exception of the elution step, the beads were washed for 5 min then placed on a magnet and the wash buffers were discarded. At the last wash the Protein low binding tubes were replaced with DNA LoBind tubes. For elution 80 μl of Elution buffer (1X m6A-IP buffer + 6.7 mM m6A nucleotides) were added directly on the beads and the tubes were incubated for 1hour with continuous shaking (1100rpm) at 4 °C. The beads were spin down and the supernatant was transferred to a clean tube. After the second round of elution the eluted RNA was precipitated using ethanol precipitation as described above. RNA pellet was resuspended in 15 μl RNase-free water and using Qubit^®^ RNA HS Assay Kit we measured the RNA concentration following manufacturer’s instructions.

#### qTNTchase-seq, qPCR, RT-PCR

RNA was metabolically labelled with BrU for 15 minutes and chased for 30 minutes as described above. RNA was purified using TRIzol following manufacturer’s instructions. 200 ug total BrU labeled RNA was used as Input for the BrU-RNA isolation. After the elution step (200 μl of 0.1% BSA and 25mM bromouridine in PBS) we added 50ul of 5X m6A-IP buffer. 4 μg (1μg ab per 500ng RNA) m6A ab were coupled to 40ul Dynabeads^®^ Protein A as described above, resuspended in 550 μl m6A-IP 1X buffer and added to the RNA mixture. RNA-antibody-beads mixture was incubated for 60 minutes at room temperature with gentle rotation. The supernatant was kept and RNA was isolated with TRIzol. The beads were washed 3 times for 5 minutes at RT (twice with low salt m6A-IP 1X buffer and last wash high salt m6A-IP 1X buffer). We eluted the RNA captured by m6A ab by competition as described in TNT-Seq section. cDNA synthesis was performed using the same amount of RNA (10–20 ng) from all fractions (Input BrU-RNA 0 min, Input BrU-RNA 30 minutes chase, Supernatant m6A-neg 0h, Supernatant m6A-neg 30 min chase, IP m6A-positive 0 min, IP m6A-positive 30 min chase). RT-PCR was performed using Q5 Hot Start High-Fidelity DNA Polymerase New England Biolabs with initial denaturation 98 °C 30s, then 32 cycles of 98 °C 10 s, 58 °C 20 s and 72 °C 55 s and final extension 72 °C 2 minutes. PCR products were resolved on agarose gel. Spike-in controls were in vitro transcribed using T7 RNA Polymerase Invitrogen following manufactures instructions. For the methylated transcripts N6-methyl-ATP (tri-link) was used in a ratio 4:1 to ATP in the in vitro transcription reaction. GFP and Luciferase sequences were used as template for the RNA transcription. For each qTNTchase-seq sample before m6A IP, *in vitro*–transcribed transcripts with and without m^6^A modification were mixed into the samples as spike-in controls at the indicated percentage of m6A-modified to m6A-unmodified transcript (Molinie et al., 2016). For all samples after BrU-IP but before m6A-IP we added 2.5×10^7^ copies from each spike included: 0% GFP, and 20% luciferase. For the *in vitro* transcribed transcripts with m6A modifications, N6-methyl-ATP (tri-link) was used in a ratio 4:1 to ATP in the *in vitro* transcription reaction. For the sequencing; Post-qTNTchase seq 1 μl of 1:2000 dilution of the universal ERCC spike-in control A (Invitrogen) was added to each fraction.

#### Transcript m6A level and splicing index

The m6A level per transcript from the qTNTchase-seq experiment were calculated as described in (Molinie et al., 2016). The ratio of the RNA abundance for each transcript between the eluate and the supernatant was represented by the ratio of the overlapping strand-specific RNA read counts normalized to the ratio of the reads of the ERCC RNAs. We used the log2-transformed read counts of ERCC RNAs to fit a linear regression model, computing the eluate ERCC reads as a function of the supernatant ERCC reads with a coefficient of 1 (Supplementary Figure 7). The log2 ratio between ERCC eluate counts and supernatant counts was indicated by the intercept of the regression formula. Only the ERCC RNAs with at least 100 read counts were used in this pipeline. M6A level = E/(E+S*2^*intercept*)

Eluate read counts (*E*), supernatant read counts (*S*), and the intercept of ERCC regression (*intercept*)

We assessed the splicing efficiency per transcript as the ratio of the overlapping strand-specific split reads (extracted by using bedtools coverage –s –F 1.0) to all (split + non-split) reads covering the transcript.

#### Quantitative real-time PCR

RNA was reverse transcribed using the Goscript reverse transcription Promega A500cDNA was quantified on an 7900HT Fast real time PCR system (Applied Biosystems) using the Go Taq qPCR Master Mix Promega (A6001). The PCR was carried out using a standard protocol with melting curve. Primers for unspliced RNA transcripts were design to span exon – intron 5’ splice junction and exon – exon boundaries for spliced RNA transcripts. Splicing efficiency was is determined by the ration of 2^-CT_spliced_ / (2^-CT_spliced_+2^-CT_unspliced_)

#### RNA sequencing and data analysis

For the BrU Chase seq, the library preparation was performed using the TrueSeq Stranded Total RNA Kit (Illumina). Sequencing was performed on an Illumina HiSeq 2500 instrument to obtain around 200M reads per sample. For the TNT-Seq, 100 ng of Input BrU-labeled fragmented RNA and 100 ng of TNT-IP eluate RNA were subjected to library preparation following the TruSeq Stranded mRNA Library Preparation Kit instructions with some modifications. The protocol started from the first strand synthesis step and 3X Clean-NA-Beads beads volume was used for the buffer exchange to include shorter RNA fragments. Mapping of strand-specific reads to GRC37 genome assembly (hg19) was done using STAR (Dobin et al., 2013) and only uniquely mapped reads were kept for further downstream analyses. To extract read coverage per nucleotide position across the genome the strand-specific bed files were sorted by chromosome and start coordinate and converted into wig files with bedtools genomecov using –scale to normalize for library size. To assess the genome-wide correlation of the m6A signal from replicates, the ratio of normalized read counts per nucleotide position of IP to Eluate, rendering the m6A signal, was converted to bigWig using wigToBigWig (UCSC) and then bigWigCorrelate (UCSC) was used. To extract the m6A signal per nucleotide position in given intervals, the depth at each nucleotide position of the examined intervals (e.g. within +/-500 bp windows around anchor points) was extracted using bedtools coverage –d –s from the m6A Input and the respective m6A IP, and then the ratio m6A IP/Input multiplied by (total number of mapped reads in the Input/ total number of mapped reads in the IP) was calculated. Then the average m6A signal was extracted at each nucleotide position from all examined entries.

#### m6A peak calling

We called m6A peaks based on a previously published pipeline (Ke et al., 2015; Ke et al., 2017). We first divided the genome into 20 bp non-overlapping bins with bedtools windowMaker and extracted the strand-specific read coverage from m6A Input and IP for all bins using bedtools coverageBed –s. Fisher’s exact test p-value was extracted from the matrix (bin Input read counts, bin IP read counts, total number of mapped reads in the Input, total number of mapped reads in the IP) and adjusted by the Benjamini and Hochberg method to determine the false discovery rate (FDR). Only windows with a p-adjusted < 0.05 in all three replicates and fold enrichment (score) minimum four in at least two out of the three replicates were kept as significant. Adjacent significant bins were merged using bedtools mergeBed into broader peaks (finally 95 % of the peaks were in the range 20–100 nt long). In the case of broad peaks, the peak summit is the midpoint of the 20 nt window with the maximum score, or the midpoint of the interval of merged adjacent bins sharing same maximum score within the same peak. In a few cases, a broad peak was assigned more than one summits if it contained non-adjacent windows sharing the same maximum score, finally yielding 58102 m6A peaks and 58311 peak summits. Custom scripts were written in awk.

#### *De novo* motif search

De novo motif search was run using HOMER (Heinz et al., 2010) within +/-150 nt intervals around the peak summit of 5651 best scoring exonic m6A peaks (minimum fold enrichment 20) and the same number of top best intronic peaks. Control sequences were generated from the respective input sequences with the scrambleFasta.pl script. Then, *de novo* motif search was run with ‘findsMotifs.pl input_sequences.fa fasta –basic –rna –len 6,7,8 –fasta scrambled_sequences’. The results were inspected in terms of enrichment, significance and the presence of common consensus sequences, with the four motifs displayed in Supplementary Figure 1B being the most represented. Those were used to scan the input sequences for the presence of match occurrences using the ‘dna-pattern’ search tool from the RSAT suite (Medina-Rivera et al., 2015) with parameters ‘search given strand only, prevent overlapping matches, origin-start, return flanking nucleotide positions 2’. Motif search was also performed in the same number of random genomic intervals as a control, generated with bedtools (–length 300 –number 5651). The matches were aligned and the logo was generated with WebLogo3 (Crooks et al., 2004).

#### Splicing kinetics and predictive models

To assess splicing efficiency we extracted the splicing index value θ as in (Mukherjee et al., 2017). θ equals to the ratio of the split reads mapping to the 5’ and 3’ SJ of an intron divided to the sum of split plus non-split reads (schematic representation in Supplementary Figure 2A). The θ value (representing Splicing Efficiency, SE) was extracted from all pulse-chase time points, for 13,532 introns with at least five reads coverage in both 5’ and 3’ SJ, and used in k-means clustering with k = 3 to call three groups of distinct splicing efficiency (fast, medium and slow) (Supplementary Figure 2B). The Splicing Efficiency Dynamics metric was calculated as SED = 1/ ((1.001-θ 0 min) * (1.001-θ 60 min)) (plotted in the *log* scale for the three groups in Supplementary Figure 2D). To assess constitutive versus alternative splicing we extracted the ψ value as in (Mukherjee et al., 2017). ψ is the ratio of constitutive split reads assigned to a given intron’s 5’ and 3’ SJ to all split reads (i.e. split reads from the given intron 5’ SJ to any downstream 3’SJ and from the intron’s 3’ SJ to any upstream 5’ SJ, as depicted in Supplementary Figure 2A). Therefore ψ is in the range 0 to 1 with 1 meaning 100 % constitutive splicing. We then used the ψ value extracted from the pulse-chase time point 60 min (closer to steady-state) to perform k-means clustering with k = 2 and define two clusters of introns, constitutive (n = 11836, minimum ψ 0.5294) and alternative (n = 1696, maximum ψ 0.5278). In the case of introns classified as alternative spliced (ψ < 0.5278) upstream or downstream exon skipping takes place.

The following features were used in logistic and linear regression models to predict splicing efficiency kinetics and alternative versus constitutive splicing:

The 5’ and 3’ splice site underlying sequence scores extracted using MaxEntScan (http://genes.mit.edu/burgelab/maxent/Xmaxentscan_scoreseq.html); distance of the 5’ SJ to the annotated transcript first start site (TSS) and of the 3’ SJ to the last end site (TES); expression calculated as coverage (reads per kb) from the m6A Input RNA-seq (15 min BrU pulse) for the whole transcript interval where the intron belongs to; intron length; intron overall m6A signal extracted as the strand-specific m6A IP read coverage divided to m6A Input read coverage, normalized by (total number of mapped m6A Input reads * total number of mapped m6A IP reads); m6A signal calculated the same way at the 5’ SJ 100 nt exonic boundary, 5’ SJ 100 nt intronic boundary, 3’ SJ 100 nt exonic boundary and 3’ SJ 100 nt intronic boundary.

To predict fast versus slow or alternative versus constitutive splicing, logistic regression was performed with R function glm (family = binomial) (all parameters apart from the sequence scores were first log scale transformed and all were then standardized). To evaluate the fitting of the model and assess discrimination, the Receiver Operating Characteristic Curve (ROC) and the area under the curve (AUC) were calculated with the R package ROCR (Sing et al., 2005). Linear regression to predict splicing efficiency using the continuous value θ (in the range 0 to 1) was performed with R function lm().

#### CLIP data analysis

We used CLIP data for SRF3 and SRSF10 from(Xiao et al., 2016)(GEO GSE71096). To calculate the relative SRSF10/SRSF3 binding per nucleotide position, we used the ModeScore column from the GEO submitted PARalyzer output file, which is the score of the highest signal divided to the sum value (signal+backround) and ranges from 0.5 to 1. We first extracted the coverage for each SRSF per nucleotide position in the +/500 nt window around 5’ or 3’ SJ, or per bin for the length-binned introns (introns with length 1000–10000 nt, binned into 1000 non-overlapping windows), by using bedtools coverage –s –d. Nucleotide positions with overlapping CLIP binding sites were assigned the cluster’s score (ModeScore column) whereas nucleotide positions with no CLIP data overlap were assigned a pseudo-score 0.1. We then computed the ratio SRFS10/SRSF3 per nucleotide position or per bin of all analyzed loci and the metagene analysis extracting the average ratio SRFS10/SRSF3 per nucleotide position or per bin was run separately for each of the subgroups fast/medium/slow or constitutive/alternative.

## ACKNOWLEDGMENTS

E.N. has been funded by a postdoctoral stipend from the Alexander von Humboldt Foundation. Work in the author’s laboratory is funded by the German Research Council (DFG) and the Alexander von Humboldt Foundation through the Sofja Kovalevskaja Award to U.A.Ø.

## Author contributions

A.L. did experimental work, interpreted data, conceived experiments, and wrote the paper. E.N. did computational analysis, interpreted data, and wrote the paper. T.C conceived experiments, interpreted data and wrote the paper. U.A.Ø. conceived experiments, interpreted data, supervised research, and wrote the paper. All authors read and approved the paper.

